# Loss of ABC transporters, White, Brown, and Scarlet, prevents increase in mitotic divisions of germline stem cells in response to mating in *Drosophila melanogaster*

**DOI:** 10.64898/2025.12.02.691910

**Authors:** Megan P. Wright, Alisa Vladimirova, Heath M. Aston, Manashree S. Malpe, Cordula Schulz

**Affiliations:** Department of Cellular Biology, University of Georgia, Athens, GA; Lake Erie College of Osteopathic Medicine, Erie, PA; Vanderbilt University Medical Center, Nashville, TN

## Abstract

The replenishment of specialized cells depends on the activity of stem cells. Recent advances in stem cell research have shown that the germline stem cells (GSCs) in *Drosophila melanogaster* can increase their mitotic activity in response to mating. Here, we show that this ability to respond to mating is eliminated if the males are mutant for either of the ABC transporters, White (W), Brown (Bw) or Scarlet (St), which are known for their role in eye pigmentation and amine production. However, reducing the expression of *w* specifically from the germline cells also caused a failure to increase GSC mitotic activity upon mating, suggesting a role of *w* in cellular fitness. The *w* gene is a common genetic background for genetic experiments and frequently used as a control. Our findings underline the importance of careful experimental design and control choice.

## INTRODUCTION

As many specialized cells, such as skin, gut, and sperm cells, are lost due to usage or death, the corresponding tissues replenish these through a process commonly referred to as tissue homeostasis [1,2]. One of the key factors in tissue homeostasis is the ability of long-term precursor cells, the stem cells, to undergo mitotic cell divisions. Their daughter cells then either become new stem cells or short-lived precursor cells that undergo a cascade of regulated proliferation and differentiation steps to replace the lost cells [3–6]. To perform tissue homeostasis effectively, stem cells must adjust their mitotic activity to situations of demand, such as injury or growth phases. Recent advances in the literature confirm that GSCs in the gonad of *Drosophila melanogaster* can adjust their mitotic index (MI^GSC^) to a variety of factors, including diet and temperature [7,8]. Specifically, male GSCs divide more frequently when the males are repeatedly mated to virgin females compared to their sibling males that are not allowed to mate [9]. How this increased demand, caused by repeated mating, is met by the GSCs is not well understood, even though some of the necessary elements have been elucidated. The increase in MI^GSC^ is dependent on the activity of G-proteins and on seven non-redundant G-protein coupled receptors (GPCRs), including three Serotonin (5HT) receptors and an Octopamine (OA) receptor [9].

In *Drosophila*, the synthesis of bioamines, 5HT, OA, and Dopamine (DA), requires the same precursor molecules as the production of eye pigments. For both pathways, these precursors are transported across membranes by the ATP-dependent ABC transporter, W [10,11]. W can form active transporters by heterodimerization with other ABC transporters, two of which have been studied in *Drosophila*, Scarlet (St) and Brown (Bw) [12–17]. In combination with St, W transports Tryptophan, which becomes synthesized into 5HT in the neurons and into the brown pigment, Ommochrome, in the eye [18,19]. Together with Bw, W transports Guanine, the precursor for Tetrahydrobiopterine (BH4). BH4 is a co-factor for enzymes that synthesize several amines, including 5HT, Dopamine, and OA in the neurons, and for the production of red eye pigments, specifically Drosopepterines [20,21].

A range of other substrates are also transported by W, including Guanosine 3’-5’ cyclic monophosphate (cGMP), Kynurenine, Riboflavin (the Vitamin B2 precursor), Xanthine, and Kynurerine [18–20,22–25]. W, but not its partners, Bw or St, is associated with cholesterol homeostasis for its role in olfactory learning. While the authors did not suggest a direct role for W in cholesterol transport, they suggested that W may form a heterodimer with one of several other unstudied ABC transporters to control the levels of cholesterol and cholesterol esters [26].

Among other tissues, expression of *w* was detected in pigmented cells, the eyes, the malphigian tubules, the testes sheath, and in non-pigmented glia and neurons of the brain [17,27,28]. The W protein was reported in vesicular fractions of several cell types, suggesting a role in intra-cellular transport. Electron-microscopy experiments revealed a localization of W in the membranes of the pigment granules within pigment cells and retinal cells, suggesting that it transports molecules from the cytoplasm into the pigment found on vesicles, and in Cos-1 cells, W was detected on the endosomal compartment [24,29]. A potential role for W in cargo influx or efflux via the plasma membrane has neither been shown, nor refuted.

Not surprisingly, animals mutant for *w* display an array of phenotypes. W mutant animals have abnormal vision, including increased sensitivity to light and abnormal phototaxis, and their retinal and dopaminergic neurons degenerate [30–34]. Along with the vision defects, *w* mutant animals display neurological phenotypes, such as learning and place memory defects and defects in locomotion [25,35–37]. *w* mutant flies also have altered sensitivity to anesthetic treatments and alcohol, and are less aggressive than *wt* flies [38–40].

Here, we show that males homozygous mutant for either of the two commonly used *w* alleles successfully mated but had no significant difference in their MI^GSC^ when compared to their non-mated siblings. In an attempt to determine how the binding partners of W factor into this effect, we found that males homozygous mutant for *bw*, *st*, or double mutant for both also had a similar MI^GSC^ as their non-mated siblings. Hence, it appears that multiple cargos passing the membranes with assistance of the ABC transporters are essential for the increase in MI^GSC^ in response to mating. Reducing *w* specifically from the germline cells resulted in the same failure to increase MI^GSC^, suggesting that cell metabolic processes limit the function of GSCs to respond to a demand for more sperm.

## RESULTS

### Animals mutant for w did not increase their MI^GSC^ in response to mating

In *Drosophila* males, GSCs are found at the tip of the testes where they attach to somatic hub cells (Figure 1A, 1A’). When one of these GSCs divide, the two daughter cells normally take on different fates. One of them becomes a new GSC that remains attached to the hub and replenishes the stem cell pool. The other becomes a gonialblast (GB) that proliferates into 16 spermatogonia. After meiosis, the cells ultimately differentiate into exactly 64 spermatids (Figure 1A)[41,42]. If a demand for more spermatids is produced via mating experiments, the GSCs divide more frequently, which can be measured by investigating the MI^GSC^ using simple immuno-fluorescence experiments [8]. The hub cells can be labeled with an antibody against FasciclinIII (FasIII) and the attached GSCs can be visualized with an antibody against Vasa. The Vasa-positive GSCs are imaged (Figure 1A’) and counted in all focal planes around the hub. By adding an antibody against phosphorylated Histone-H3 (pHH3), cells in mitosis become apparent and the percentage of the GCSs that are actively dividing, the MI^GSC^, can be calculated [8].

**Figure 1.**
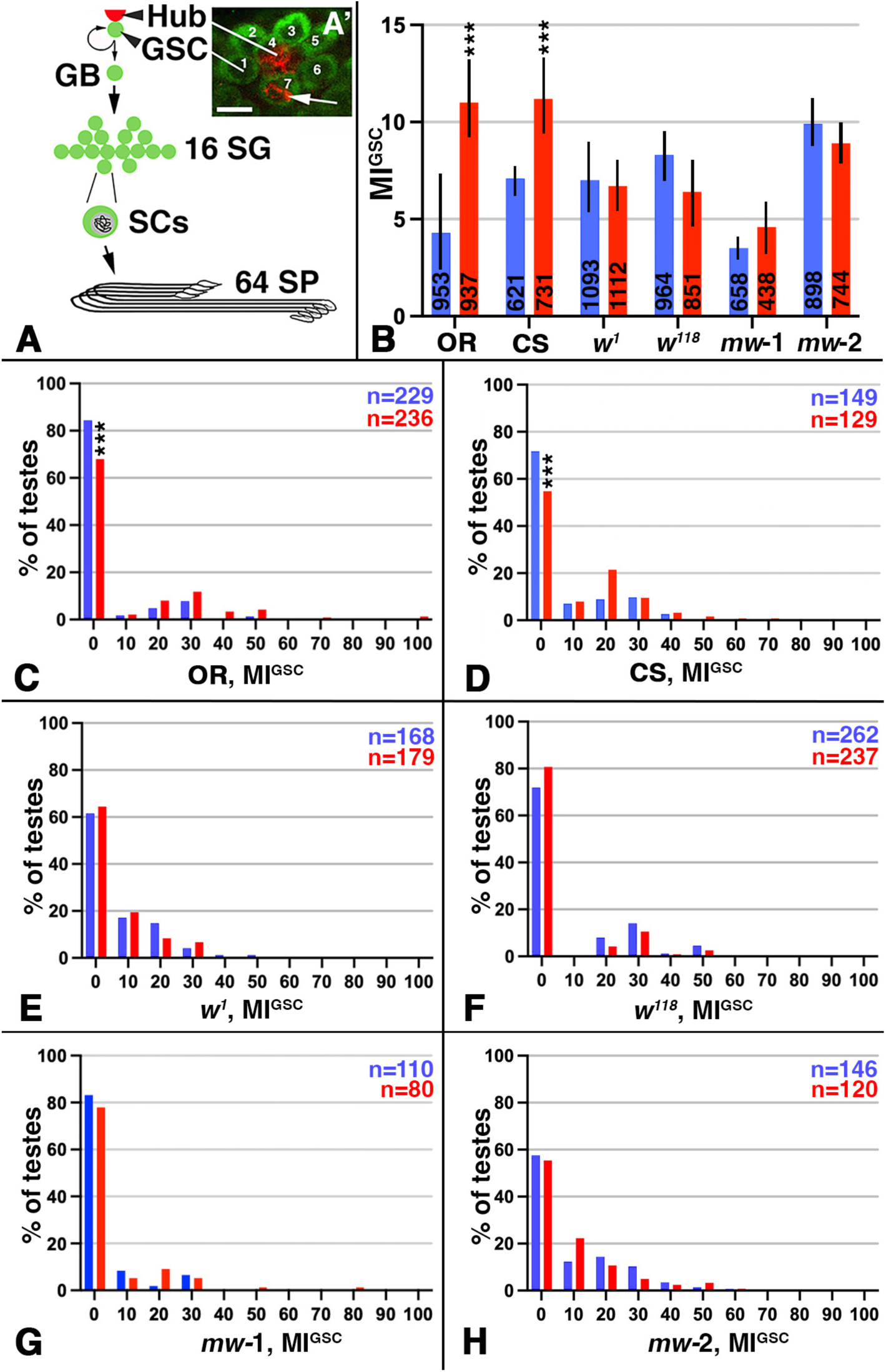
Mated *w* mutant males failed to increase MI^GSC^. A) Cartoon depicting how a GSC division results in a new GSC and a stem cell daughter that will ultimately produce 64 spermatids (SPs). GB: gonialblast, SG: spermatogonia, SC: spermatocyte. A’) Immuno-staining of the apical tip of a *wt* testis, showing seven Vasa-positive GSCs (green) next to the FasIII-positive hub (red). One of the seven GSCs is in mitosis based on anti-pHH3-staining (red; arrow). Scale bar: 10μm. B-H) Blue: non-mated condition, red: mated condition, ***: P-value < 0.001, numbers of GSCs in bar graphs and number of gonads (n=) in FDGs as indicated. B) Bar graph showing MI^GSC^ of wt, *w* mutant, and *mw* males as indicted. C-H) FDGs showing median of bin of MI^GSC^ across populations of males on the X-axes (bin width=10) and the percentage of testes with each MI^GSC^ on the Y-axes; genotypes as indicated.

While *wild-type* (*wt)* males always significantly increase MI^GSC^ in response to repeated mating, we noted considerable inconsistencies in our data when males were kept in a *w* mutant background. Intrigued by this observation, we set out to investigate if the failure of males to increase MI^GSC^ might be due to the *w* mutation itself. We employed two of the commonly used *w* alleles, *w^1118^* and *w^1^*, and set up single male mating experiments with these, as well as with two *wt* strains, *Oregon R* (*OR)* and *Canton S (CS)*. To obtain reproducible data, we always use 100 males for the mated group and another 100 males for the non-mated controls for each experiment. To illustrate the overall MI^GSC^, we add the GSC numbers and calculate MI^GSC^ from three replicate experiments and display the differences between non-mated and mated animals in bar graphs. These show that *OR* and *CS* males clearly increased MI^GSC^ when repeatedly mated, and that animals mutant for *w^1118^* or *w^1^*displayed similar MI^GSC^ under both conditions (Figure 1B).

A GSC division is not a frequent event. Most animals within one group, or population, have zero GSCs in division. To assure that the differences in MI^GSC^ between non-mated and mated males are not due to a few unusually behaving males, but by an overall MI^GSC^ increase within the group of males, we use Frequency Distribution Graphs (FDGs). FDGs group the males based on different MI^GSC^ and then display how often those MI^GSC^ are represented among the testes of one group of males. In the FDGs, the *wt* mated males had a significant decrease in the number of GSCs with a MI^GSC^ of zero and showed more testes with MI^GSC^ above zero (Figure 1C, D). For *w* mutant males, in contrast, we did not observe a reduction in the number of GSCs with a MI^GSC^ of zero and did not see more testes with MI^GSC^ above zero upon mating (Figure 1E, F). All groups of males used in our experiments mated with virgin females based on visual observations and the production of progeny (Table 1). We conclude that *w* is required for the increase in MI^GSC^ in response to mating.

**Table 1.** Fertility Assay. Male fertility was calculated based on the % of females that produced offspring after mating with males of the indicated genotype on day one of the experiment. BL#: Bloomington stock number.

| Genotype | BL # | Male fertility |
| --- | --- | --- |
| *OR* x *OR* | N/A | 72% |
| *CS* x *CS* | N/A | 62% |
| *w*<sup>1118</sup> x *w*<sup>1118</sup> | N/A | 81% |
| *w*<sup>1</sup> x *w*<sup>1</sup> | 679 x 679 | 70% |
| *bw*<sup>1</sup> x *bw*<sup>2b</sup> | 245 x 248 | 79% |
| *mw*-1 | N/A | 69% |
| *mw*-2 | N/A | 74% |
| NG4-1 | N/A | 85% |
| NG4-2 | 4937 | 87% |
| *bw*<sup>1</sup> x *bw*<sup>1</sup>*st*<sup>1</sup> | 245 x 686 | 64% |
| *bw*<sup>2b</sup> x *bw*<sup>1</sup>*st*<sup>1</sup> | 248 x 686 | 70% |
| *st*<sup>1</sup> x *bw*<sup>1</sup>*st*<sup>1</sup> | 605 x 686 | 77% |
| *bw*<sup>1</sup>*st*<sup>1</sup> x *bw*<sup>1</sup>*st*<sup>1</sup> | 686 x 686 | 50% |
| W33641xNG4-1 | 33641 | 76% |
| W33644xNG4-1 | 33644 | 86% |

### The *mini-white* (*mw*) gene does not restore the ability of *w* males to respond to mating

*mw* is a version of *w* that lacks the first intron and is commonly used as a reporter to demonstrate the presence of a mobile element in the fly genome [27]. Expression of m*w* can restore many defects seen in *w* mutants. For example, expression of *mw* rescued the eye pigmentation and retinal degeneration defects in *w* animals [25]. However, the expression levels of *w* and *mw* underlies position-variegation and the insertion in different locations of the genome produce flies with varying eye colors, that range from light yellow to orange to red [43–45] The *mw* reporter also contains an insulator that can severely reduce its expression levels [46]. To overcome the limitation of expression levels, more than one copy of *mw* is often used for rescue experiments. For example, four copies of the *mw* gene were used to rescue the defects of *w* mutant males in copulation success [47].

We attempted to rescue the effect that mutations in *w* have on MI^GSC^. For this, we used two fly lines that carry four copies of the *mw* gene (*mw*-1 and *mw*-2, Table 2). These are the same fly lines we use for our mating experiments below, but in a *w* mutant background. As expected, expression of *mw* in these lines restored eye color to a dark red (data not shown). However, it did not restore testis sheath pigmentation defects (data not shown), probably because the *mw*-constructs lacks the necessary regulatory elements [27]. When we performed single male mating experiments with either *mw* line, we also did not see a rescue in the response to mating of the *w* mutant males (Figure 1B). The FDG created from this mating trial showed no significant decrease in the number of testes at an MI^GSC^ of zero and no increase in testes with an MI^GSC^ above zero (Figure 1G, H). Since we did not succeed in restoring the defects in *w* mutants via *mw* expression, we performed the following experiments with flies that had been crossed into a *wt* X-chromosomal background (referred to as NG4 flies).

**Table 2:** Transgenic flies used in this study. Abbreviation and complete genotypes of fly stocks are shown

| Abbreviation | Genotype |
| --- | --- |
| mw1 | w-/Y; UAS-dicer (mw+)/UAS-dicer (mw+); nanos-Gal4-VP16 (mw+)/nanos-gal4-VP16 (mw+) |
| mw2 | w-/Y; UAS-dicer (mw+)/UAS-dicer (mw+); nanos-Gal4 (mw+)/nanos-gal4 (mw+) |
| NG4-1 | X/Y; UAS-dicer (mw+)/UAS-dicer (mw+); nanos-Gal4-VP16 (mw+)/nanos-gal4-VP16 (mw+) |
| NG4-2 | X/Y; UAS-dicer (mw+)/UAS-dicer (mw+); nanos-Gal4 (mw+)/nanos-gal4 (mw+) |
| w<sup>33641</sup> | y, v<sup>1</sup>; P{TRiP.HMS00042}attP2/TM3, Sb<sup>1</sup> |
| w<sup>33644</sup> | y, v<sup>1</sup>; P{TRiP.HMS000425}attP2 |

### The bw and st genes were also required for the increase in MI^GSC^ in mated males

Since Bw and St are both W’s binding partners, we hoped to pinpoint the activity of W in the response to mating to a specific pathway. Hence, we conducted our mating experiments with *bw* and *st* single, and double mutant males, as listed in Figure 2. As obvious from the bar graph, control males (*bw^1^, st^1^/wt*) showed the expected increase in MI^GSC^, while all the mutants failed to respond to mating (Figure 2A). Likewise, the FDG of the mutants howed no significant decrease in the number of testes at an MI^GSC^ of zero and no increase in testes with an MI^GSC^ above zero (Figures 2B-H), We conclude that transport via the Bw and St transporters is also essential for the normal GSC behavior in mated males.

**Figure 2.**
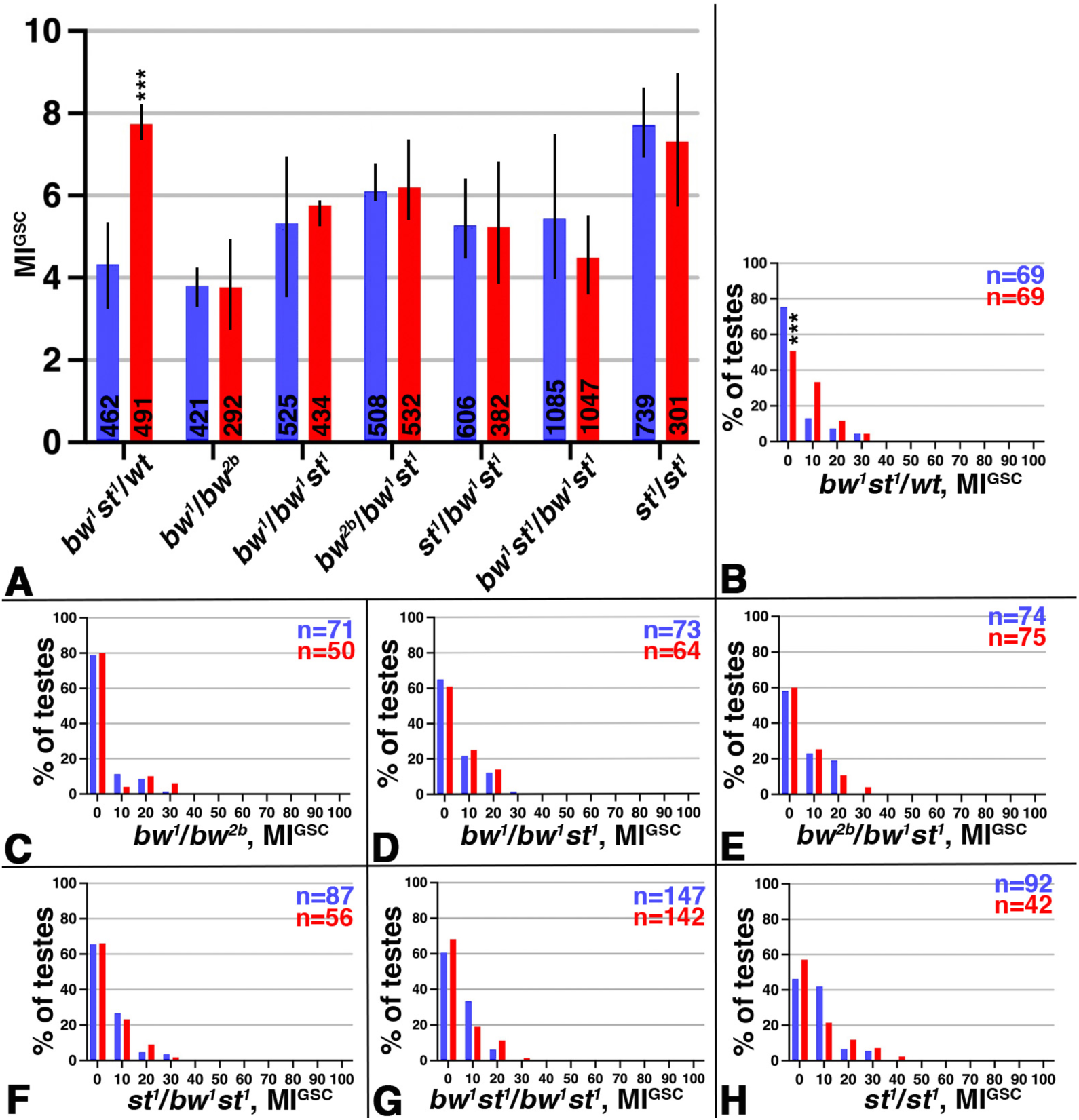
Mated *bw* and *st* males failed to increase MI^GSC^. A-H) Blue: non-mated condition, red: mated condition, ***: P-value < 0.001, numbers of GSCs in bar graphs and number of gonads (n=) in FDGs as indicated. A) Bar graph showing MI^GSC^ of *wt*, *bw*, and *st* mutants and double mutants, as indicted. B-H) FDGs showing median of bin of MI^GSC^ across populations of males on the X-axes (bin width=10) and the percentage of testes with each MI^GSC^ on the Y-axes; genotypes as indicated.

### Reducing w from the germline reproducibly blocked the increase in MI^GSC^ in response to mating

Wondering if the W transporter is dispensible within the germline cells for the increase in MI^GSC^ in response to mating, we used tissue-specific knock-down of *w* via two independent RNA-interference (RNAi) lines (w^33641^ and w^33644^)[48,49]. To achieve reduction specifically within the germline, we crossed the RNAi flies to the commonly used germline-Gal4-transactivator, NG4 [50]. The two NG4 lines (NG4-1 and NG4-2, Table 2) had previously been successfully used for mating experiments [9]. Progeny from *wt* animals crossed to either of these NG4 or to either of the RNAi lines showed the stereotypical increase in MI^GSC^, but when either of the two *w* RNAi lines were crossed to either of the NG4, their progeny failed to increase MI^GSC^ in response to mating (Figure 3A). Consistent with this, the FDG for the control crosses showed a significant decrease in the number of testes from mated males with a MI^GSC^ of zero and an increase in the number of testes with an MI^GSC^ above zero (Figure 3B, C), while the FDGs for the *w* RNAi crosses did not show it (Figure 3D,E).

**Figure 3.**
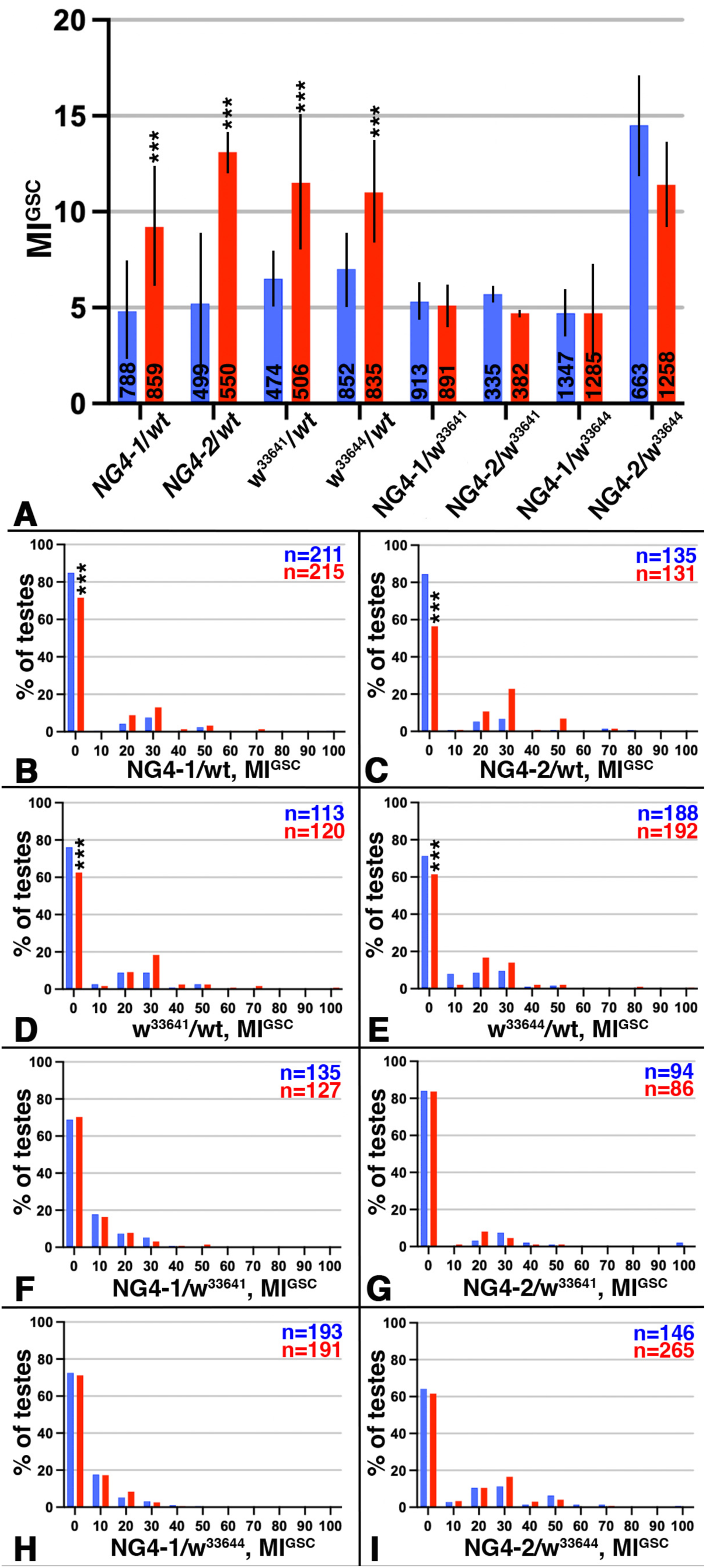
Decreasing the expression of *w* in the germline via RNAi repeatedly lead to a failure to increase MI^GSC^ after mating. A-I) Blue: non-mated condition, red: mated condition, ***: P-value < 0.001, numbers of GSCs in bar graphs and number of gonads (n=) in FDGs as indicated A) Bar graph showing MI^GSC^ of control (*wt* crossed to either NG4 or RNAi line) and experimental (either NG4 crossed to RNAi lines), as indicted. B-I) FDGs showing median of bin of MI^GSC^ across populations of males on the X-axes (bin width=10) and the percentage of testes with each MI^GSC^ on the Y-axes; genotypes as indicated.

## DISCUSSION

The *w* mutation is heavily used as a marker in genetic and molecular studies, but the underlying cellular and biochemical functions of *w* are normally not considered as contributing factors to a phenotype. Yet a variety of abnormal behaviors have been associated with the *w* mutation. Here we show that loss of the W transporter from whole males or reduction specifically in the male germline through RNAi blocked the increase in MI^GSC^ after mating. This failure to respond to mating could not be rescued by expressing several copies of the *mw* gene. A reasonable explanation for this observation is that the *mw* constructs lack a gonad-specific enhancer element or that we could not reach high enough expression levels from the *mw* genes used in this study.

GSCs are neither pigmented cells, nor neurons, suggesting a different function for W within the germline. It is possible that W interacts with an unidentified ABC transporter in the germline to specifically regulate mitotic divisions. However, the pleiotropic effect of *w* on the flies and the variety of substances transported by the W protein suggests that the failure of the mutant males to increase their MI^GSC^ could have multiple underlying reasons.

Interestingly, *w* affects mating behavior. Expression of *w* throughout the male flies from a heat-shock driven transgene, or loss of *w* function, caused increased male sexual arousal and male to male courtship behavior, even in the presence of females [29,51–53]. Loss of *w* also drastically slows copulation success of virgin males, but not so much of mating-experienced males [47,53]. Though we cannot exclude the possibility that loss of *w* has an effect on the fly mating behavior in our experiments, we doubt this is directly linked to their failure to increase MI^GSC^. We performed all experiments with single males (which excludes male to male courtship) and the males were non-virgins, excluding a reduced copulation success. This is supported by the results of our fertility tests that show that *w* mutant males copulated successfully with an similarly large number of virgin females as *wt* males did (Table 1).

The effects of the *w* mutation outside the eye have been associated with a role for *w* in the nervous system, mediated by 5HT and DA. For example, 5HT is essential for place memory and plays a role in phototaxis, and DA was associated with male aggression [34,36,39]. Consistent with a role for neurotransmitters in *w* phenotypes, several previous studies have shown that *w* mutant animals have abnormal levels of amines, including DA and 5HT [28,36,54]. Mutants for the two known binding partners of W, Bw and St, also contained abnormal amine levels [28]. Together with our observation that these mutants also failed to increase MI^GSC^ upon mating, it opens the possibility that the increase in MI^GSC^ in mated *wt* males is dependent on amines from the nervous system. This appears consistent with our previous findings that amine receptors are required for the increase in MI^GSC^ in response to mating [9].

In addition to a possible role for w in the nervous system in regulating MI^GSC^, we show that loss of *w* from the germline cells also impaired the ability of the mated males to respond to mating. During aging, the intestinal stem cells of *Drosophila* increase the expression of *w*, as well as the frequency of their divisions. Interestingly, loss of *w* reduced intestinal cell proliferation due to a role in metabolism that *mw* did not restore [55]. We did not detect a significant difference in the expression of *w* in testes tips from mated compared to non-mated *wt* males [56]. However, we agree with the idea of a role for *w* in stem cell metabolism. Likely, any mutation, genetic situation, or environmental factor could reduce the fitness of the GSCs, and, hence, interfere with their performance, such as decreasing the frequency of mitotic divisions. We also showed that at least seven GPCRs are required for increasing MI^GSC^ in response to mating [9]. It is tempting to speculate that some of the ligands for these receptors may not be instructive, but rather permissive due to a role in the fitness of the cells.

Overall, we found that the increase in MI^GSC^ after mating is dependent on W, Bw, and St, suggesting that they play some role in regulating GSC divisions, even if this role is potentially permissive. These findings highlight the importance of careful planning of mating experiments, which should all be performed in a *wt* X-chromosomal background.

## MATERIALS AND METHODS

### Fly husbandry

Flies were raised on a cornmeal/agar diet and kept in incubators with temperature-, light-, and humidity-control. The *OR*, *CS*, NG4-1, NG4-2, and *w^1118^*flies were originally obtained from the Bloomington stock center but have been in the laboratory for more than 10 years, while the *w^1^*, *bw*, and *st* alleles and the RNAi-flies were obtained more recently (Tables 1+2)[57,58]. *X*⌃*X*, *y,w,f* /Y/*shi^ts^* flies were provided by Barry Ganetzky.

### Mating experiments

Mating experiments were performed at 29°C. Flies were fed with yeast paste on apple juice-agar plates in egg lay containers for 24 hours prior to the experiment. Males were placed into one slot of a mating chamber either by themselves (non-mated) or with three virgin females (mated). The chambers were closed by apple juice-agar lids supplemented with yeast paste. On each of the following two days, females were discarded, and each mated male was provided with three new virgin females. Apple juice-agar lids supplemented with yeast paste were replaced daily for both conditions. Female virgins from the stock *X*⌃*X*, *y,w,f*/Y/*shi^ts^* were used for mating experiments.

### Immuno-fluorescence and microscopy

Animals were placed on ice to immobilize them. Gonads were dissected in Tissue Isolation Buffer (TIB) and immediately placed into a 1.5 ml tube with cold TIB buffer. Gonads were then fixed, stained and imaged as previously described [8,59]. The mouse anti-FasciclinIII (FasIII) antibody (1:10) developed by C. Goodman was obtained from the Developmental Studies Hybridoma Bank, created by the NICHD of the NIH and maintained at The University of Iowa, Department of Biology, Iowa City, IA 52242. The goat anti-Vasa antibody (1:50-1:500) was purchased from Santa Cruz Biotechnology Inc. (sc26877). The rabbit anti-Vasa antibody (1:3000) was a gift from Ruth Lehmann. The rabbit anti-phosphorylated Histone H3 (pHH3) antibodies (1:100 to 1:1000) were purchased from Fisher (PA5-17869), Millipore (06-570), and Santa Cruz Biotechnology Inc. (sc8656-R). Secondary antibodies coupled to Alexa 488, 568, and 647 (1:1000) and Slow Fade Gold/DAPI embedding medium were purchased from Life Technologies. Staining was observed with a Zeiss Axiophot, and images taken with a digital camera and an apotome via the Axiovision Rel. software.

### Graphic presentations and data analysis

Bar graphs and FDGs were generated with GraphPad prism version 10.2.3 and assembled into composite images using Adobe Photoshop. The GraphPad prism default two-tailed student’s t-test was used to analyze statistical relevance.

## ACKNOWLEDGEMENTS

The authors are grateful to Leon McSwain, Alicia Hudson, Chederli Gaile Balbuen Belongilot, and Kyona Garrett for technical assistance. We thank Bruce Baker for helpful discussions and Maya Barfield for help with figure generation. We are grateful to Barry Ganetzky for the *X*⌃*X, y,w,f/Y/shi^ts^* fly stock, to Ruth Lehmann for the rabbit anti-Vasa antibody, and to Wolfgang Lukowitz for the use of his microscope. This work was supported by NSF grants #0841419 and #1355009.

